# LNP-CpG: deploy the self-adjuvant role of mRNA vaccines

**DOI:** 10.64898/2026.08.19.745633

**Authors:** Ning Luan, Han Cao, Xiaolong Zhang, Fengmei Yang, Caixia Lu, Yang He, Qianqian Li, Yanwei Bi, Zhanlong He, Shengtao Fan, Liu Liu, Shaogui Wan, Cunbao Liu

## Abstract

With the rapid advancement of mRNA vaccines, lipid nanoparticles (LNPs) have emerged as pivotal carriers and adjuvants for non-mRNA vaccine modalities, driven by their superior nucleic acid delivery efficiency and intrinsic self-adjuvanting properties. In this study, we systematically evaluated various formulation strategies combining LNPs and the CpG adjuvant within a varicella-zoster virus glycoprotein E (VZV-gE) subunit vaccine framework. We demonstrated that uniform nanoparticles formed by LNP-encapsulated CpG (LNP-CpG), when simply admixed with the gE antigen, elicited superior immunogenicity compared to alternative encapsulation configurations. Intramuscular administration of a two-dose (LNP-CpG)+gE regimen significantly augmented both humoral and cellular immune responses in mice, markedly outperforming the commercial vaccine Shingrix (administered at a 1/10 human dose). Crucially, the identical regimen induced robust, comparable immune profiles to a full human dose of Shingrix in rhesus macaques. Furthermore, LNP-CpG displayed broad-spectrum utility across diverse vaccine platforms, demonstrating efficacy against both respiratory (RSV) and neurotropic (HSV) pathogens, compatibility with multiple modalities, including subunit (VZV-gE, RSV- Pre-F), live-attenuated (LA-HSV), and inactivated (i-HSV) vaccines; and versatile implementation in a combined VZV+RSV formulation. Collectively, our findings position LNP-CpG as a versatile, safe, highly efficacious adjuvant platform with substantial clinical translational potential, offering a compelling paradigm for next-generation vaccine development.

## Introduction

Messenger RNA (mRNA) therapeutics have emerged as a powerful class of medicines with demonstrated clinical benefit in infectious disease, cancer and genetic disorders^1, 2^. Lipid nanoparticles (LNPs), the most effective nonviral delivery vehicles for mRNA, can elevate the cellular immune response, have attracted great attention due to unique advantages such as simple formulation, good biocompatibility, and large payload^3, 4, 5^. Beyond mRNA, LNPs serve as a versatile vaccine adjuvant delivery system (VADS) for delivering other nucleic acids, peptides, and immunomodulatory molecules^6^. Crucially, LNPs function not merely as passive delivery vehicles; they exhibit intrinsic self-adjuvating properties, allowing them to be utilized as standalone vaccine adjuvants for the protein subunit vaccine platform^7, 8, 9^. By leveraging a membrane fusion mechanism, LNPs precisely internalize their encapsulated payloads into antigen-presenting cells (APCs), thereby enhancing delivery and translation efficiencies while promoting cross-presentation of antigens.

LNPs are composed of four primary lipid components: an ionizable cationic lipid, a helper phospholipid, cholesterol, and a PEGylated (PEG, polyethylene glycol) lipid^10, 11^. By rapid mixing, those four components can form uniform and stable particles. The ionization of the cationic lipid affects the surface charge of the lipid nanoparticle under different pH conditions. This charge state can influence plasma protein absorption, blood clearance and tissue distribution as well as the ability to form endosomolytic non-bilayer structures^12^. Ionizable lipid, which is responsible for the self-adjuvating properties of LNP, contains positively charged ionizable amine groups (at low pH) to interact with the anionic nucleic acid, during particle formation and also facilitate membrane fusion during internalization^13^. PEGylated lipid is a critical component that governs the size and homogeneity of LNPs, and act as a steric barrier to prevent aggregation during storage^14^. Concurrently, the particle size of LNPs significantly influences their downstream immunogenicity and in vivo trafficking. The average particle sizes of LNP candidates are generally maintained 60-100 nm. Specifically, smaller particles drain more easily to lymph nodes and are preferentially engulfed by APCs, while larger particles are more readily phagocytosed by macrophages^15,16^.

Synthetic CpG oligodeoxynucleotides (CpG-ODNs, briefly called CpG) are short single-stranded DNA molecules containing unmethylated CpG dinucleotides and a phosphorothioate backbone^17^. CpG 1018, a B type CpG, has been approved by the FDA for use in Heplisav-B, a hepatitis B vaccine. CpGs are potent Toll-like receptor 9 (TLR9) agonists, which strictly located within the endosomes of dendritic cells (DCs), and then triggering a TLR9-Myd88 pathway, further induce the upregulation of proinflammatory cytokine and type I interferon (IFN) genes in macrophages, DCs, and B cells^18^. The maturation of DCs upregulating the expression of co-stimulatory molecules, and further attracting a larger pool of T cells, such as effector T cells (CD4^+^) and cytotoxic T lymphocytes (CD8^+^). Compared with traditional alum adjuvants, CpG motifs can robustly activate a Th1-biased cellular immune response^19, 20, 21, 22^.The combination of CpG and alum can effectively reverse the inherent Th2-biased immune response of alum alone, significantly augmenting cellular immunity while maintaining its robust capacity for high-titer antibody induction. The encapsulation of CpG within particles or lipid-based components protects it from nuclease-mediated degradation, thereby further augmenting its immunogenicity and driving a more potent immune response, such as PLGA, VLP, and nanomaterials ^23, 24, 25^.

Several studies have demonstrated the potent immunogenicity of CpG encapsulated within LNPs, with such lipid-based adjuvants proving highly effective in vaccines targeting viral infections and tumors^26, 27, 28^. In this study, we systematically optimized the formulation and preparation of CpG-loaded LNPs for Varicella-zoster virus (VZV) subunit vaccines, which inherently necessitate a robust cellular immune response. Our findings revealed that encapsulating CpG alone within LNPs (LNP-CpG), followed by the external addition of the gE antigen, elicited immune responses either comparable or superior to those induced by Shingrix^TM^ in both mice and rhesus macaques. The prepared LNP-CpG adjuvant formulation exhibited excellent stability, and its ’mix-and-use’ strategy with external antigens facilitates broad-spectrum applicability across diverse pathogen vaccine designs. Consequently, we extended the deployment of this adjuvant platform to subunit vaccines for RSV and combined RSV+VZV, as well as to alternative modalities such as attenuated and inactivated HSV vaccines. Across these diverse pathogens and vaccine platforms, the LNP-CpG adjuvant consistently demonstrated a powerful capacity to drive immune responses, significantly elevating antigen-specific antibody titers, neutralizing antibody levels, and cell-mediated immunity. Furthermore, vaccines formulated with this novel adjuvant provided superior protective efficacy in viral challenge models.

## Results

### 1. LNP-CpG adjuvanted VZV subunit vaccine induces robust humoral and cellular immunity in mice

A precise optimization of the ionizable cationic lipid: PEGylated lipid ratio is essential for tailoring the physicochemical properties and therapeutic performance of LNP-based formulations. In this study, an officially disclosed LNP benchmark formulation of 50:10:38.5:1.5 from mRNA-1273^10^ (Moderna) is used for our VZV subunit vaccine platform. In our previous studies, we thoroughly explored the adaptation of these LNP compositions for VZV vaccines^29, 30, 31, 32^.

The class B CpG ODN1018, further denoted as CpG, or VZV antigen gE, were encapsulated in LNP by rapid mixing. Four different LNP formulations were generated (Fig.1a): gE were encapsulated alone-LNP-gE; CpG admixed in LNP-gE-(LNP-gE)+CpG; CpG were encapsulated alone, then gE admixed -(LNP-CpG)+gE; gE and CpG were all encapsulated in LNP-LNP-(CpG+gE). Dynamic light scattering (DLS) analysis (Fig.1b) indicated all LNPs has uniform and low polydispersity index (PDI) (<0.3), but (LNP- CpG)+gE has the smallest diameter of 100.3 nm. In this group, LNP has CpG encapsulation efficiency of 95.6%, equals 14.34 μg/dose CpG (Fig.1c).

After two doses of immunization with 28 days interval, immunized serum was collected for detection of gE-specific IgG titers (Fig.1d). Non-adjuvanted gE induced negligible IgG titers, as expected. With CpG adjuvanted, IgG titers elevated 10-fold in group gE+CpG than gE-alone. Group (LNP-CpG)+gE induced the highest level of humoral immunity, with IgG geometric mean titer (GMT) of 298,667, two folds higher than Shingrix^TM^ 1/10 hd (120,000) (Fig.1e). In the test of cellular immunity, group (LNP-CpG)+gE also induced the higher Th1-biased cytokine secretion, including IL-2 and IFN-γ, with tests of flow cytometry (FC), enzyme-linked immunosorbent assay (ELISA), and enzyme-linked immunospot assay (ELISpot) of gE-stimulated splenocytes (Fig.1f-h).

We further investigate the structural properties of LNP-CpG adjuvant. Cryo-transmission electron microscopy (cryo-TEM) image revealed that LNP-CpG exhibit polydisperse nanoscopic spherical structures comprising “bleb” structures with distinctly different electron density (Fig.1i). Differences in morphology between individual LNP, including bleb formation has recently been attributed to occur during buffer exchange^33^, and such bleb formation can improve cellular endocytosis efficiency of immuno-stimulator. Using the identical formulation, fluorescently labeled LNP-CpG was prepared and subsequently formulated with IRDye 800CW-labeled gE antigen- (^Did^LNP- ^Cy3^CpG)+^800CW^gE (Fig.S1). ICR mice were administered a single dose via intramuscular injection, followed by continuous monitoring of *in vivo* fluorescence intensity at predetermined time points (10h, day1, day2, day3, day4, day8) as well as the *ex vivo* biodistribution of fluorescent signals in isolated tissues and organs. In vivo imaging revealed a progressive decline in fluorescent signals at the local injection site over time. Intriguingly, however, *ex vivo* organ imaging demonstrated that both ^Did^LNP and ^800CW^gE exhibited massive accumulation within the intestinal tissue at 10 hours post-injection (Fig.S1c-d). Over the subsequent days (from day 1 to day 2), this intestinal fluorescence gradually diminished, accompanied by a synchronous, delayed signal accumulation in the liver. This sequential trafficking profile indicates that following intramuscular administration, the LNP-CpG formulations preferentially drain into the local lymph nodes and subsequently aggregate in the abdominal cisterna chyli, which is the anatomical region boasting the densest immune cell population and richest lymphatic network in the body, before eventually entering the systemic bloodstream to circulate through the liver.

### 2. LNP-CpG adjuvanted VZV subunit vaccine induces robust innate immunity in rhesus macaques

Nonhuman primate serves as an important animal model for preclinical testing of vaccines due to their genetic and physiological similarities to humans. Having fabricated the vaccine, we compared (LNP-CpG)+gE’s efficacy with Shingrix^TM^ human dose (hd) and Ganwei^TM^ hd for induction of both humoral and cellular immune responses in rhesus macaques. Since only two VZV vaccines are currently approved in China: Shingrix^TM^ from GSK, a subunit vaccine adjuvanted by AS01b, recommended two doses for full schedule; and Ganwei™, an Oka-based live attenuated vaccine developed by Changchun BCHT Biotechnology, officially recommended one dose.

As illustrated in Fig. 2a, rhesus macaques in the (LNP-CpG)+gE and Shingrix™ hd groups were administered two intramuscular (I.M.) doses on day 0 and 28. In contrast, the Ganwei™ hd group received a single subcutaneous (S.C.) dose on day 0. On day 42, whole anticoagulated blood was collected from all animals for the isolation of peripheral blood mononuclear cells (PBMCs) (Fig. 2a). In (LNP-CpG)+gE group, 500 μL (LNP- CpG)+gE were injected, comprising of 143 μg CpG and 50 μg gE antigen, equals ten-mice- doses (also equals hd). We first measured the early innate immune responses after vaccine administration by monitoring the fluctuation of distinct leukocyte subsets. Both subunit vaccine groups (Shingrix^TM^ hd and (LNP-CpG)+gE) were shown similarity to induce an increase trend in circulating leukocytes at 14 days after second dose immunization (Fig. S2b). Analysis of the individual dynamics of various inflammatory cell types revealed that this elevation was predominantly driven by three granulocyte subsets: neutrophils, basophils, and eosinophils (Fig. S2a). These results demonstrate the long-term and sustained stimulatory effect of the vaccine on the immune system, indicating that systemic immunity remains continuously activated even 14 days post-boost, which is a well-documented phenomenon in vaccine immunization^34, 35^.

**Figure 1.**
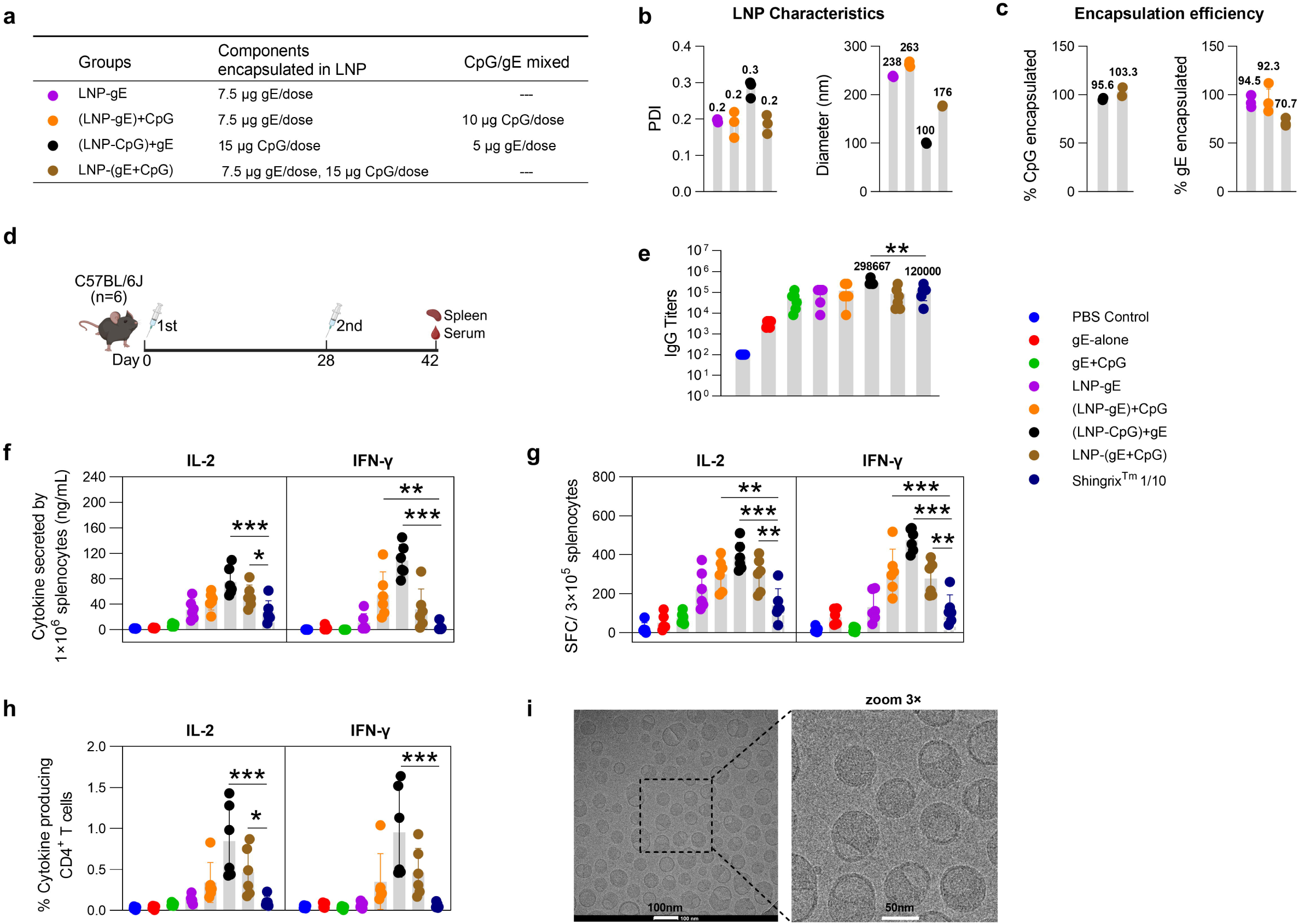
LNP characterization and immunity induced by VZV subunit vaccines in mice. (a) Components encapsulated or mix immediately in LNPs. (b) DLS analysis of PDI and Z-average diameter. (c) Encapsulation efficiency of CpG and gE were measured by OliGreen and BCA assay, respectively (*n* = 3, mean ± SD). (d) immunization schedule. (e) gE specific IgG titers of immunized serum after two-dose vaccines injected (day 42). (f-h) The recall responses of IL-2 and IFN-γ in splenocytes were evaluated using ELISA (f), ELISpot (g), and flow cytometry assays (h). Data in (e-h) from vaccine groups were analyzed by one-way ANOVA (using a Dunnett’s multiple comparisons test, compare the mean of each column with the mean of group Shingrix^TM^ 1/10). Data were shown as mean ±SD. *, *p* ≤ 0.05; **, *p* ≤ 0.01; ***, *p* ≤ 0.001. (i) Cryo-transmission electron microscopy (cyro-TEM) image of LNP-CpG. Scale bar represents 100 nm and 50 nm.

**Figure 2.**
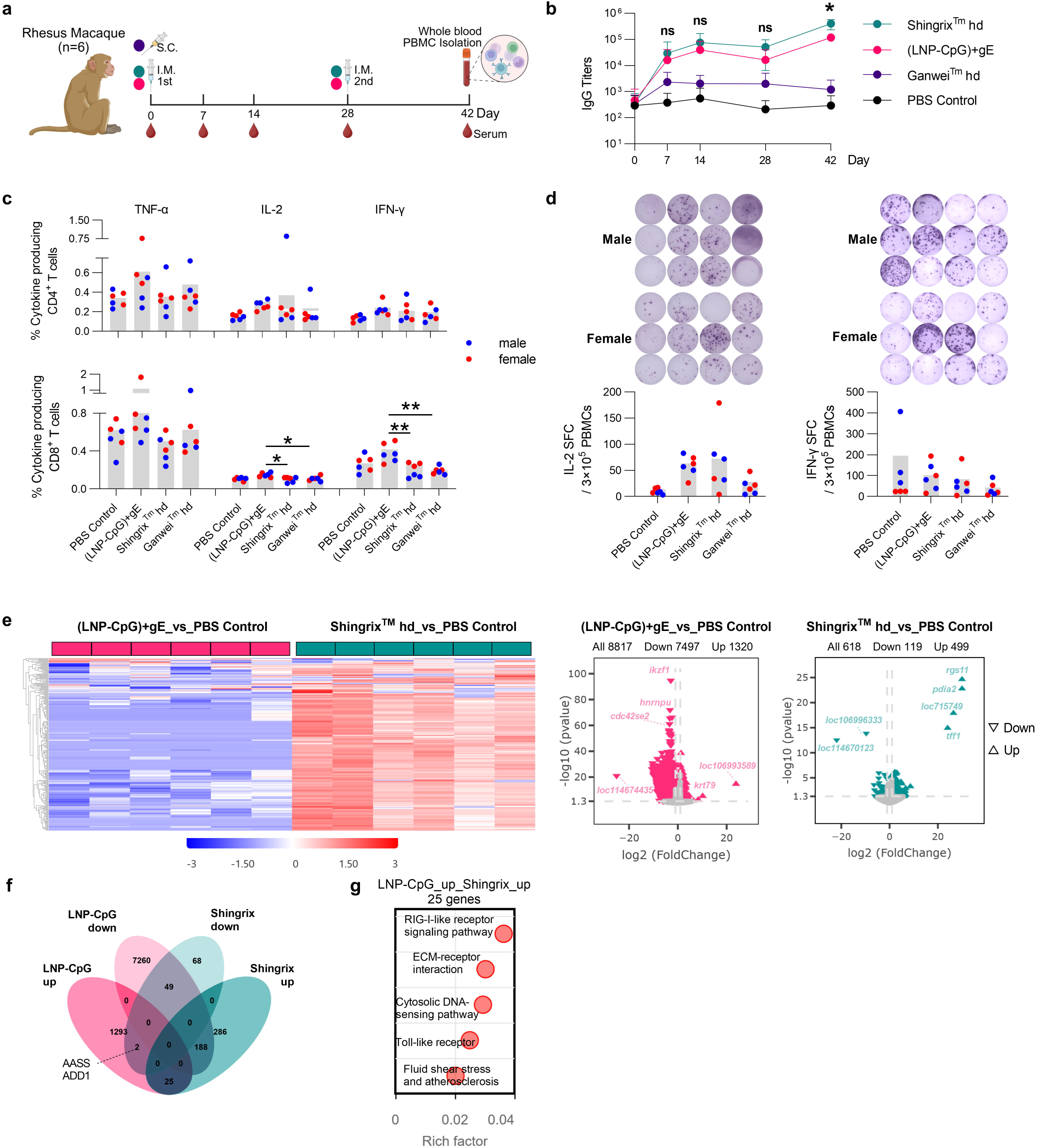
Immunological profiling and comparative transcriptomic analysis of the (LNP- CpG)+gE VZV subunit vaccine in rhesus macaques. (a) Immunization schedule. Rhesus macaques (*n* = 6 per group) were immunized subcutaneously (S.C.) with a single dose of Ganwei™, or intramuscularly (I.M.) twice at a 28-day interval (Days 0 and 28) with Shingrix™ hd) or (LNP-CpG)+gE. Whole blood was collected and PBMCs were isolated 14 days after the final immunization for subsequent analyses. (b) Kinetic changes in gE- specific IgG antibody titers following immunization. Statistical significance between the Shingrix™ hd and (LNP-CpG)+gE groups at each time point was determined using a two-way ANOVA with Bonferroni’s multiple comparisons test (*, *p* ≤ 0.05). (c) Proportions of TNF-α-, IL-2-, and IFN-γ-producing cells among CD4^+^ and CD8^+^ T cells analyzed by flow cytometry. (d) Quantification of IL-2- and IFN-γ-secreting cells per 3×10^5^ PBMCs assayed by ELISpot, with representative images shown in the top panels. In (c-d), results for male (blue dots) and female (red dots) animals are presented separately to evaluate the potential impact of sex on experimental outcomes. (e) Heatmap and volcano plot analyses of differentially expressed genes (DEGs) in the Shingrix™ hd and (LNP-CpG)+gE groups relative to the PBS control group. (f) Venn diagram of overlapping DEGs among upregulated and downregulated gene sets from the LNP-CpG and Shingrix™ groups. (g) Bubble plot showing the top enriched KEGG pathways for the 25 co-upregulated DEGs shared between the LNP-CpG and Shingrix™ groups. Data are presented as mean ±SD.

Blood samples were collected on day 0, 7, 14, 28 and 42. Shingrix^TM^ hd group always keep the highest IgG titers, reach to 10^4^ on day 14 and 10^6^ on day 42. gE-specific IgG titers induced by (LNP-CpG)+gE were similarly robust and in the same order of magnitude as those elicited by Shingrix^TM^ hd. Ganwei^TM^ hd group reach the lowest level of around 10^3^ in all test time points (Fig. 2b). PBMC was isolated from the whole blood at day 42, cellular immune responses were assayed by FC and ELISpot (Fig. 2c-d). Th1 cytokines- tumor necrosis factor (TNF-α), interleukin-2 (IL-2), interferon gamma (IFN-γ)-were elevated significantly in (LNP-CpG)+gE group compared with other commercial vaccine groups.

Besides, the safety profile among vaccine groups were also been assessed. Animal rectal temperatures were continuously monitored within 72 hours following both the first and second vaccinations. After the first dose, the temperature trends in all vaccine groups showed no significant difference compared with the PBS control group (Fig. S3a). Following the second dose, however, the (LNP-CpG)+gE and Shingrix^TM^ hd groups exhibited a transient temperature elevation within the first 32 hours post-injection, which subsequently returned to baseline levels; no notable differences were observed at other time points relative to the PBS control (Fig. S3b). Similarly, no abnormal changes in body weight were observed after either vaccination, with all vaccine groups maintained a comparable trend to the PBS control (Fig. S3c). For hematological and biochemical evaluations, anticoagulated blood samples were collected prior to vaccination (day 0), as well as after the first dose (day 28) and the second dose (day 42). These whole-blood samples were sequentially analyzed using an automated hematology analyzer to determine coagulation profiles, biochemical parameters, and complete blood counts (CBC). Regarding the coagulation profile (Fig. S3d), the variation trends in the (LNP-CpG)+gE group were highly consistent with those in the PBS control group, remaining well within the normal physiological range and indicating no risk of hemorrhage. For serum biochemical analysis, eight core parameters were evaluated to monitor organ functions: TBIL (μmol/L), ALT (U/L), AST (U/L), and ALP (U/L) for hepatic function (Fig. S3e); CREA (μmol/L) and BUN (mmol/L) for renal function (Fig. S3f); and TC (mmol/L) and LDH (U/L) for metabolic assessment (Fig. S3g). Summarily, (LNP-CpG)+gE formulation demonstrated an excellent in vivo safety profile, with no significant alterations observed in key hematological and biochemical parameters (coagulation, hepatic, renal, and metabolic panels) compared to the PBS control and commercial vaccines.

To gain a more in-depth understanding of the innate immune activation, whole blood gene expression changes elicited by the two kinds of VZV subunit vaccines were determined by transcriptomic analyses on PBMCs of monkey 24 h post 1^st^ vaccine injection. Heatmap analysis of differentially expressed genes (DEGs) (false discovery rate-adjusted p-value≤0.01, │log_2_FoldChange│≥5) revealed that, compared with the PBS control group, immunized with LNP-CpG adjuvanted VZV subunit vaccines in rhesus macaques induced a predominant gene downregulation profile, whereas Shingrix^TM^ hd exhibited a greater degree of gene upregulation. Volcano plot analysis further demonstrated that the LNP- CpG group induced 1,320 upregulated and 7,497 downregulated DEGs, whereas the Shingrix group induced 499 upregulated and 119 downregulated DEGs (Fig. 2e).

The overlap between significantly DEGs for each group was determined (Fig. 2f), identifying two genes that were uniquely upregulated in the LNP-CpG group but downregulated in the Shingrix^TM^ hd group: *AASS* and *ADD1*, therefore likely representing the VZV vaccine specific response. Previous studies indicate that *AASS* is closely associated with mitochondrial function and redox homeostasis, whereas *ADD1* plays crucial roles in maintaining cell membrane stability, regulating Na⁺/K⁺-ATPase activity, and participating in signal transduction and ion transport^36, 37, 38^. Pathway enrichment analysis of the 25 co-upregulated DEGs shared by both LNP-CpG and Shingrix^TM^ hd groups showed that the top five KEGG pathways ranked by rich factor were: RIG-I-like receptor signaling pathway, ECM-receptor interaction, cytosolic DNA-sensing pathway, Toll-like receptor signaling pathway, and fluid shear stress and atherosclerosis (Fig. 2g).

### 3. LNP-CpG modulate innate immunity in TLR9 pathway

Since the promising ability of inducing robust cellular and humoral immune responses in VZV-gE subunit vaccines, as demonstrated by rodent and non-human primates. The mechanism of action of this vaccine type was further investigated in detail. After gE delivered to DCs by LNP-CpG, it’s important for sufficient antigen escape from lysosome to be presented. Previous report has showed that cytokine production by bone marrow cells in response to LNP-CpG is completely dependent on TLR9^28^, but failed to localize LNP-CpG in cell. Herein, we use confocal microscopy to observe the localization of LNP- CpG in matured BMDC. In the absence of LNP-CpG (0 µg/mL), the gE antigen showed high colocalization with lysosomes, indicating that the FITC-labeled antigen was internalized by Lyso-Tracker-labeled lysosomes (Fig.3a). In the merged confocal results, with 0.5 μg/mL LNP-CpG treated, BMDCs shows lower pearson’s correlation coefficient than untreated, indicates more FITC-gE escape from the lysosome in this group (Fig.3b).

To systematically delineate the intracellular trafficking and antigen-processing pathways of the LNP-CpG formulation, a panel of pharmacological inhibitors was implemented (Fig. 3c). Cells were pretreated with cytochalasin D (CD, an actin polymerization inhibitor) to scrutinize the dependency of cellular uptake on macropinocytosis. To further investigate downstream post-internalization fates, bafilomycin A1 (BA1) or concanamycin B (CB), two vacuolar H^+^-ATPase inhibitors, were employed to arrest endosomal acidification, thereby examining whether the ionizable lipid-mediated endosomal escape of CpG is strictly pH-dependent. Through confocal microscopy observation and pearson’s correlation coefficient analysis, we found that LNP-CpG- mediated antigen escape from lysosomes is largely driven by endosomal acidification. Specifically, the addition of the CD inhibitor did not affect antigen escape, whereas treatment with BA1 or CB significantly reduced the extent of escape, restoring the Pearson correlation coefficient to approximately 0.6 (Fig. 3d). Additionally, to evaluate the contribution of the proteasome pathway to subsequent gE antigen processing and cross-presentation, epoxomicin (a highly selective proteasome inhibitor) was introduced to perturb cytoplasmic antigen degradation. However, due to severe drug-induced cytotoxicity, no valid data were obtained.

**Figure 3.**
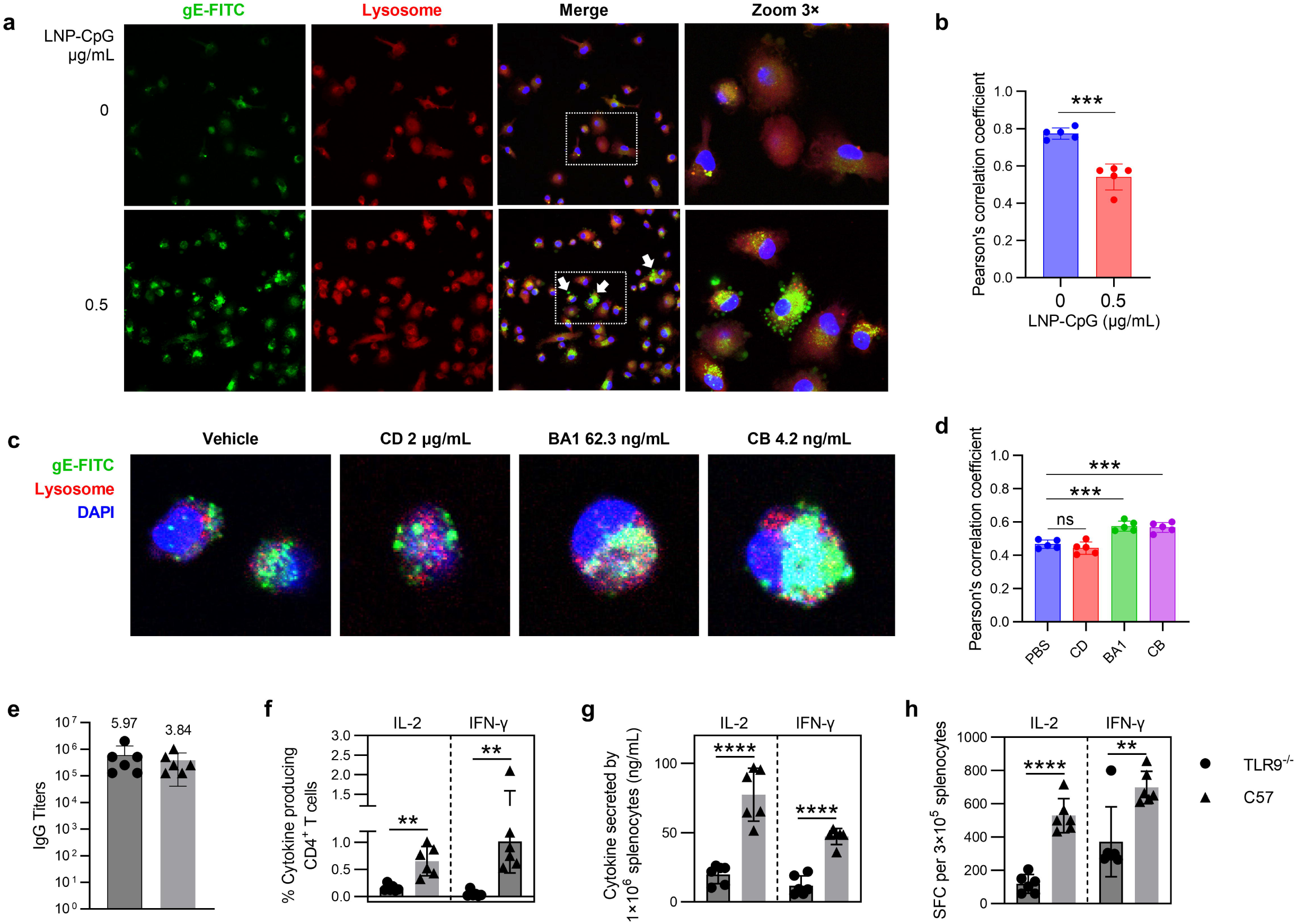
Characterization and immunostimulatory evaluation of LNP-CpG in BMDCs and TLR9^-/-^ mice. (a) Confocal microscopy images depicting the colocalization of FITC-labeled gE antigen with lysosomes (labeled by LysoTracker) in BMDCs stimulated with LNP-CpG (0 or 0.5 µg/mL). Nuclei of cells were counterstained with DAPI (1 µg/mL) for 5– 10 min at room temperature. (b) Pearson’s correlation coefficient analysis quantifying the colocalization in (a). (c) Confocal microscopy images showing the colocalization of FITC-labeled gE antigen with LysoTracker-labeled lysosomes in BMDCs pretreated with various inhibitors, followed by stimulation with 0.5 µg/mL LNP-CpG and FITC-gE antigen. (d) Pearson’s correlation coefficient analysis quantifying the colocalization in (c). (e–h) Cellular and humoral immune responses in TLR9^-/-^ and wild-type (C57BL/6) mice immunized with a two-dose regimen of the gE subunit vaccine adjuvanted with LNP-CpG: (e) Serum IgG antibody titers determined by ELISA. Geometric mean titers (GMTs, ×10⁵) are indicated above the symbols for clarity. (f– h) Antigen-stimulated splenocyte immune responses evaluated by flow cytometry (f), ELISA (g), and ELISpot (h) assays. In (b) and (f-h), data were analyzed by unpaired *t* test (two tailed). In (d), data were analyzed by one-way ANOVA (using a Dunnett’s multiple comparisons test, compare the mean of each column with the mean of PBS column). **, *p* ≤ 0.01; ***, *p* ≤ 0.001; ****, *p* ≤ 0.0001. CD, cytochalasin D; BA1, bafilomycin A1; CB, concanamycin B.

As we known, CpG ODN is localized to the endosomes or lysosomes after cellular internalization and binds to endosomal TLR9. TLR9^-/-^ mice (a C57bL/6J background) were used to identify the immune response of LNP-CpG adjuvanted VZV-gE vaccine. In the humoral immunity, it seems that TLR9^-/-^ mice did not display a declined IgG titer, but rather increased IgG with no significant difference (Fig. 3e). In the cellular immune response, TLR9^-/-^ mice suggested the most impaired immunity, decreased about 5-fold compared with naïve C57 mice in CD4+ T cell FC test, 3-fold in ELISA and ELISpot test (Fig. 3f-h).

## 4. LNP-CpG exhibit high protectivity in respiratory virus subunit vaccines

Based on the fact that a simple mixture of LNP-CpG and pathogen antigens is sufficient to elicit its characteristic activation of innate immune pathways, we applied this adjuvant formulation to subunit vaccines for various other pathogens.

First, in the respiratory syncytial virus (RSV) subunit vaccine, we mixed the LNP-CpG adjuvant (80 μL/dose) with the pre-F antigen (2 μg/dose). Naive Balb/C mice were administered two intramuscular (I.M.) doses of RSV subunit vaccines with 0-28 day; whole blood was collected on days 49 (three weeks after final immunization) (Fig. 4a). Pre-F and post-F specific IgG titers in immunized serum were evaluated by ELISA (Fig. 4b). It was observed that LNP-CpG group induce significantly superior IgG titers than Alum group, whether specific to pre-F and post-F. Compared to the Alum group, the LNP-CpG group demonstrated an approximately 6-fold increase in the titers of pre-F-specific IgG antibodies. Furthermore, in contrast to the Alum group, which induced a Th2-biased response (with an IgG2a/IgG1 ratio of 0.03, far less than 1), the LNP-CpG group significantly induced a Th1- biased immune response (with an IgG2a/IgG1 ratio of 3.22, far greater than 1). The neutralizing antibody titers (nAbs) of the immune sera against RSV A2 and RSV B strains were also determined based on the number of infection plaques in Hep-2 cells (Fig. 4c). Similarly, the LNP-CpG adjuvant was able to induce nAbs that were significantly higher than those induced by the Alum group. After 4 dpi, mice were sacrificed and lung were dissected, RSV viral loads in the lungs of mice were determined by qPCR (Fig. 4d); the (LNP-CpG)+Pre-F group exhibited lower pulmonary viral loads compared to the Alum+Pre- F group. These results demonstrate that when the LNP-CpG adjuvant is utilized in an RSV subunit vaccine, it elicits stronger immunogenicity and provides a higher level of protection in mice than the Alum adjuvant.

**Figure 4.**
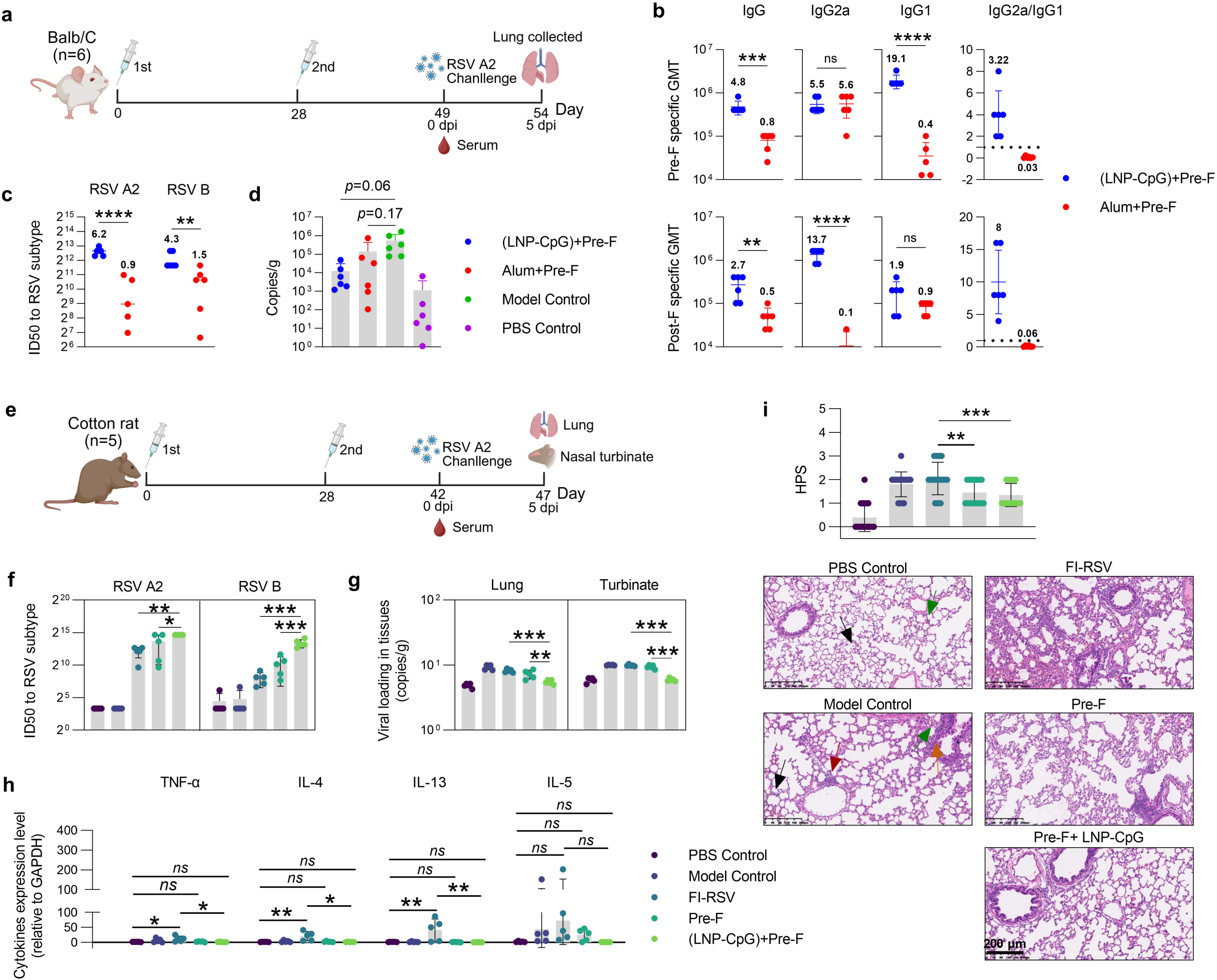
Immune profiling of (LNP-CpG)+Pre-F RSV subunit vaccine in mice and cotton rat. (a) Schematic delineating the vaccine administration protocol, RSV A2 challenging, and data collection timeline for Balb/C mice (*n* = 6). (b) Immune sera were collected after two doses injection of vaccine, pre-F and post-F specific IgG, IgG2a, and IgG1 antibody production as evidenced by ELISA assay. Geometric mean titers (GMTs, ×10⁵) are indicated above the symbols for clarity. The IgG2a/IgG1 ratio was calculated using individual values, with the mean values also indicated. (c) Neutralizing antibody titers (nAbs) against RSV A2 and RSV B strains were determined by an IC50 assay using HEp- 2 cells. In (b) and (c), data from group (LNP-CpG)+Pre-F and group Alum+Pre-F were analyzed by unpaired t test (two tailed). (d) Viral loads (copies/g) of RSV A2 in lung tissues (5 dpi) were quantified by qPCR. Data from group (LNP-CpG)+Pre-F, Alum+Pre-F, and model control were analyzed by one-way ANOVA (using a Dunnett’s multiple comparisons test with model control as a control column). (e) Schematic delineating the vaccine administration protocol, RSV A2 challenging, and data collection timeline for Cotton rat (*n*=5). (f) nAbs of immune serum from cotton mice against RSV A2 and RSV B strains were also determined by IC_50_ assay. (g) Viral loads (copies/g) of RSV A2 in lungs and nasal turbinates (5 dpi) were quantified by qPCR. In (f) and (g), data from group FI-RSV, Pre-F, and (LNP-CpG)+Pre-F were analyzed by one-way ANOVA (using a Dunnett’s multiple comparisons test, compare the mean of each column with the mean of every other column). (h) Expression level of cytokines (TNF-α, IL-4, IL-13, IL-5) in lung tissue from challenged cotton rat were determined by qPCR. Data were analyzed by one-way ANOVA. (i) Histopathological analysis and representative micrographs of left lung tissues post- infection. Lung sections were stained with H&E, and histopathological scores (HPS) were independently assigned by four blind reviewers. Moderate to severe inflammatory cell infiltration observed in the alveoli, perivascular region, bronchioles, and interstitium were indicated by black, red, orange and green arrow, respectively. Data from model control and vaccine groups were analyzed by one-way ANOVA (with correction for multiple comparisons using a Tukey test). *, *p* ≤ 0.05; **, *p* ≤ 0.01; ***, *p* ≤ 0.001; ****, *p* ≤ 0.0001. Illustration in (a) and (e) created in BioRender.

Comparatively, cotton rats are more susceptible to RSV than Balb/c mice and are recognized as an better animal model for RSV infection. Previous report has shown formalin-inactivated RSV vaccines, FI-RSV, exhibit high risk of enhanced respiratory disease (ERD) and was halted consequently^39, 40^. We evaluated the efficacy of FI-RSV, and subunit vaccines, including Pre-F only and (LNP-CpG)+Pre-F in cotton rats. Similarly, cotton rats received two I.M. vaccine injections, and whole blood was collected 14 days after the final immunization to obtain immune serum (Fig. 4e). From the results of nAbs, for both RSV A2 and RSV B, the (LNP-CpG)+Pre-F vaccine group exhibited the highest titer values (Fig. 4f). Correspondingly, this group also demonstrated the lowest pathogen loads in the lungs and nasal turbinates following RSV A2 challenge (Fig. 4g). In this cotton rat model, FI-RSV—unsurprisingly—exhibited the lowest nAbs and the highest levels of pulmonary inflammatory cytokines (Fig. 4h). Furthermore, hematoxylin-eosin (H&E) staining results revealed that FI-RSV induced extensive focal accumulation of inflammatory cells within the lungs (Fig. 4i). In contrast, the subunit vaccine groups—particularly the (LNP-CpG)+Pre-F group—significantly reduced the accumulation of pulmonary inflammatory cytokines, including TNF-α, IL-4, IL-13, and IL-5. Correspondingly, histopathological sections of the lungs from this group demonstrated only mild lesions, thereby demonstrating the exceptional protective efficacy of the (LNP-CpG)+Pre-F vaccine in the cotton rat model.

Furthermore, the LNP-CpG adjuvant was successfully utilized to formulate a combination RSV+VZV subunit vaccine containing Pre-F and gE (Figure S4a). Compared with traditional aluminum adjuvants, the LNP-CpG-adjuvanted combination vaccine demonstrated significantly elevated Pre-F- and gE-specific IgG titers, enhanced nAb titers, and robust cell-mediated immune (CMI) responses, while providing superior protection against body weight loss and mortality in challenge models. Compared to the single Pre-F antigen formulation, the LNP-CpG adjuvant combined with both Pre-F and gE antigens slightly increased Pre-F-specific IgG titers (with no statistically significant difference; Figure S4b), not significantly decreased nAbs to RSV A2 and RSV B (Figure S4c), and significantly enhanced IFN-γ-related CMI responses as measured by ELISA, ELISpot, and FC (Figure S4d). Conversely, when compared with the single gE antigen formulation, the combination formulation reduced gE-specific IgG titers approximately three-fold and partially attenuated CMI responses, particularly IL-2-related responses (Figure S4e). Following live RSV A2 challenge on day 49, daily body weight monitoring over 5 days showed that mice in the LNP-CpG groups experienced only transient weight loss on day 1 before rapidly recovering, maintaining better overall body weight compared to the unimmunized model control group (Figure S4f). Quantification of viral load in lung tissues at 5 dpi (Figure S4g) revealed that Pre-F-containing LNP-CpG formulations (both monovalent and combination) substantially reduced lung viral titers compared to the model control group (*p* < 0.001), demonstrating robust viral clearance and effective protection against viral challenge in vivo. Overall, the LNP-CpG adjuvant shows promising prospects for research and application in combined VZV and RSV vaccines. This finding aligns with our previous conclusions from the RSV+VZV combined mRNA vaccine study^41^. However, the optimal loading capacity of LNP-CpG for external antigens requires further exploration to ensure that the efficacy of the combined subunit vaccine reaches a level comparable to that of individual subunit vaccines.

### 5. LNP-CpG can significantly enhance the protective efficacy of HSV vaccines, particularly inactivated HSV vaccines

We continued to apply the LNP-CpG adjuvant to traditional herpes simplex virus (HSV) vaccines, including live attenuated HSV-1 vaccine (LA-HSV) and inactivated HSV-1 vaccine (i-HSV). A prime-boost I.M. immunization strategy (day 0 and 28) was applied to Balb/c mice. To comprehensively evaluate HSV vaccine efficacy, animals were randomly allocated into non-challenged and challenged cohorts. The non-challenged cohort was utilized exclusively to characterize vaccine-elicited immune profiling. Subsequently, the challenged cohort was longitudinal monitored for body weight fluctuations, infection symptom scores, survival rates, and vaginal viral shedding. At 7 dpi, vagina tissues, spinal cords, and dorsal root ganglia (DRG) were collected for viral load quantification, serving as key metrics to assess protective immunity (Fig. 5a).

**Figure 5.**
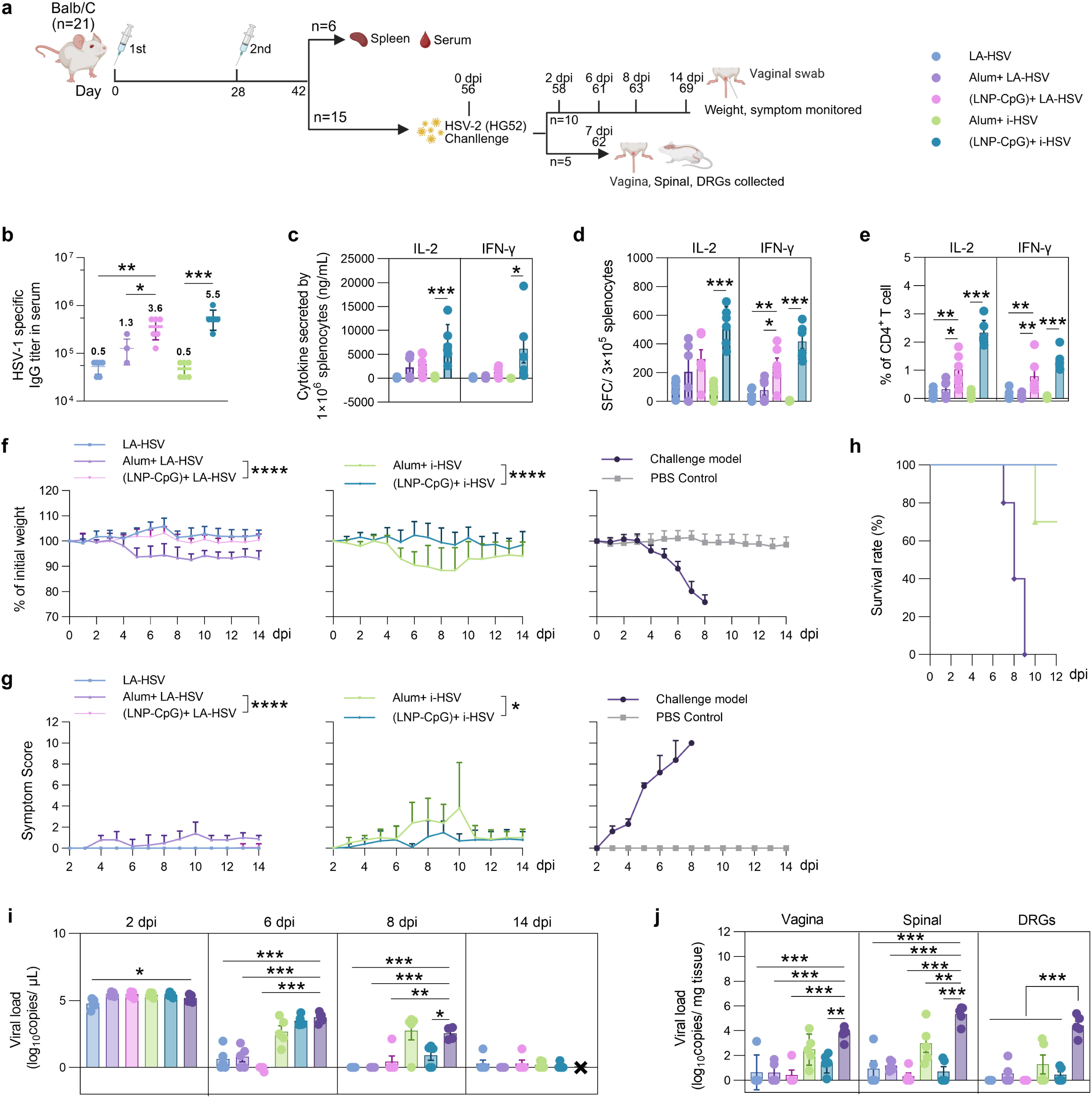
Immune profiling of live attenuated and inactivated HSV vaccines in mice (a) Schematic representation of the immunization schedule, HSV-2 challenging, and data collection timeline for Balb/C mice (*n* = 21). (b) immune sera were collected 14 days after two doses injection of vaccine, HSV-1 specific IgG titers were detected by ELISA (*n* = 6). Geometric mean titers (GMTs, ×10⁵) are indicated above the symbols for clarity. (c) The concentration of IL-2 and IFN-γ cytokines released by 1×10^6^ splenocytes was measured by ELISA (ng/mL). (d) The frequencies of IL-2- and IFN-γ-secreting cell spots per 3×10^5^ splenocytes were quantified via ELISpot assay. (e) The percentages of CD4^+^IL-2^+^ and CD4^+^IFN-γ^+^ T cells within the1×10^6^ splenocytes population were determined by FC. Data were analyzed by one-way ANOVA (with correction for multiple comparisons using a sidak test). After challenging, 10 mice were chosen to monitor the weight loss (f), infection symptom score (g), and survival rate (h). In (f) and (g), data from group Alum and LNP- CpG adjuvanted groups were analyzed by unpaired t test (two tailed). (i) At intermediate time points (2, 6, 8, and 14 dpi), vaginal swabs were collected from six randomly selected mice per group for viral load quantification via qPCR. ’X’ in the last column indicates missing data due to animal mortality in the Model Control group by day 9. (j) Five mice were sacrificed at 7 dpi, vagina, spinal and DRGs were collected, viral load was determined by qPCR. In (i) and (j), data were analyzed by one-way ANOVA (using a Dunnett’s multiple comparisons test with model control as a control column). *, *p* ≤ 0.05; **, *p* ≤ 0.01; ***, *p* ≤ 0.001; ****, *p* ≤ 0.0001. Illustration in (a) created in BioRender.

Compared with alum, LNP-CpG induced higher IgG titers and CMI response in both LA-HSV and i-HSV vaccines. It’s notably that LNP-CpG adjuvant demonstrated a more pronounced enhancement of both humoral and cellular immune responses for the i-HSV than for LA-HSV. Specifically, LNP-CpG boosted IgG antibody titers by 2.8-fold (LA-HSV) and 11-fold (i-HSV) relative to the conventional alum adjuvant (Fig. 5b). Regarding cellular immune responses, ELISA, ELISpot, and flow cytometry assays consistently revealed the same trend: LNP-CpG elicited a several-fold to dozens-of-fold increase over the aluminum adjuvant in the LA-HSV, whereas it triggered a hundred-fold to thousand-fold elevation in the i-HSV (Fig. 5c-e).

Following the HSV-2 strain HG52 challenge, the model control group displayed progressively worsening symptoms of infection (Fig. 5g), with weight loss initiating at 4 dpi (Fig. 5f). Mortality onset occurred at 6 dpi, culminating in a 100% mortality rate by 9 dpi (Fig. 5h). Conversely, except for the Alum+i-HSV group which incurred 3 deaths at 10 dpi, all other vaccine formulations demonstrated robust protective efficacy, successfully preventing severe post-challenge weight loss and alleviating clinical symptoms. Notably, despite its suboptimal performance in generating immune responses, the unadjuvanted LA-HSV formulation still conferred 100% protection against the HG52 challenge, significantly outperforming the Alum-adjuvanted group (Alum+ LA-HSV). Viral load analysis from vaginal swabs sampled at scheduled intervals during infection revealed that all vaccine formulations significantly reduced viral shedding, with the sole exception of the Alum+ iHSV group (Fig. 5i). Furthermore, LA-HSV formulations, with or without adjuvant, decreased viral loads more pronouncedly than the i-HSV group, which tightly mirrored the body weight and clinical symptom data. An identical trend was observed in the viral load quantification of vagina tissue, spinal cords, and DRGs harvested at 7 dpi (Fig. 5j).

As observed from the viral load results in Fig.5i and 5j, the incorporation of the LNP- CpG adjuvant significantly enhances the protective efficacy of HSV vaccines, regardless of the antigen platform (LA-HSV or i-HSV). Specifically, supplementing the LA-HSV vaccine with the LNP-CpG adjuvant ((LNP-CpG)+LA-HSV) facilitated an earlier reduction in viral loads within vaginal swabs (<6 dpi) (Fig. 5i). Conversely, while the addition of LNP- CpG to the i-HSV vaccine ((LNP-CpG)+i-HSV) also succeeded in lowering viral loads, its protective onset was delayed, becoming progressively pronounced only at 7 dpi (Fig. 5j). Taken together, the integration of the LNP-CpG adjuvant into both HSV vaccine modalities effectively elicits superior immune response levels and confers robust protective outcomes.

## Discussion

Although vaccines that induce protection through antibodies are routinely used in clinical practice, there remains a need for vaccines that elicit potent and sustained T-cell immunity for specific infections. As a neurotropic and lymphotropic virus, VZV establishes latency in the nervous system upon primary infection and can reactivate when immunity declines, leading to herpes zoster (HZ). CMI induced by VZV vaccines, rather than humoral immunity, plays a decisive role in defending against VZV reactivation and controlling the severity of HZ.

Shingrix^TM^, the only licensed VZV subunit vaccine, holds a significant advantage in market vaccination due to its strong humoral and cellular immune responses. However, AS01B adjuvant component remains a major concern on Shingrix^TM^, resulting from the production capacity and high reactogenicity of QS-21. In this study, LNP-CpG adjuvanted VZV subunit vaccine, comparable, even stronger efficacy were induced in mice (Fig.1) and rhesus macaques (Fig.2), but low reactogenicity were observed (Fig.S3). VZV-specific T-cell response is a critical confederation for controlling latent reactivation^42^, both CD4^+^ and CD8^+^ T cell responses have been shown to be induced at higher levels by LNP-CpG adjuvanted VZV vaccine than other two licensed VZV vaccines.

Although traditional liposomes—such as the DOPC/cholesterol-based vehicle in AS01B—can mimic cell membrane structures to deliver immunomodulatory agents like MPL and QS21, they struggle to efficiently deliver pathogen-associated molecular patterns (PAMPs) that require recognition by intracellular pattern recognition receptors (PRRs). As a specific agonist for intracellular TLR9, CpG requires an effective lipid-based carrier for intracellular delivery. Encapsulating CpG in such carriers not only protects it from nuclease degradation to boost utilization efficiency, but also effectively activates intracellular TLR9, thereby eliciting a robust cellular immune response. Considerable efforts, including our own and those of other groups ^25, 28, 43^, have focused on encapsulating CpG within diverse nanoparticle platforms. For instance, the encapsulation of CpG2395 and BW006 within poly(lactic-co-glycolic acid) (PLGA) nanoparticles yielded an encapsulation efficiency of less than 30%^25^. Although LNPs co-encapsulating CpG2395, BW006, and the gE antigen matched the immunogenicity of Shingrix™, they exhibited an undesirably large particle size and suboptimal encapsulation efficiencies (70% for CpG and 60% for gE), resulting in significant raw material waste^29^. Furthermore, while LNPs have been employed to deliver D-type CpG in influenza split vaccine development, they failed to demonstrate a clear advantage over the conventional alum adjuvant in inducing higher IgG titers or Th1 responses^9, 44^. Collectively, these findings confirm the feasibility of LNP-CpG as a vaccine adjuvant; however, the lipid formulation of LNPs and their compatibility with CpG and antigens warrant further optimization.

In this study, we aimed to identify the optimal assembly platform for combining antigens and CpG in a VZV subunit vaccine designed to elicit robust CMI responses. We discovered that encapsulating CpG alone within lipid nanoparticles (LNP-CpG) and subsequently mixing them with the gE antigen yielded an ideal particle size of approximately 100 nm (Fig.1b). Through optimization of our lipid composition, the encapsulation efficiency of CpG within the LNPs exceeded 95% (Fig.1c). Furthermore, this external antigen-loading strategy effectively prevented gE antigen loss during the encapsulation process. Notably, LNPs within the 100–150 nm diameter range have been reported to exhibit superior cellular uptake, immunogenicity, and safety *in vivo*^45^. Presenting the antigen on the exterior of adjuvant particles promotes its recognition and activation by APCs, thereby facilitating the targeted delivery of CpG into APCs. Given the inherent capability of LNPs to deliver nucleic acids (e.g., mRNA, siRNA) into APCs and facilitate their release from phagosomes or endosomes into the cytoplasm, our experimental results demonstrated that the LNP-CpG adjuvant likewise promotes endosomal-to-cytosolic antigen release through a lysosomal acidification-dependent mechanism (Figure 3a-d).

Although this lipid nanoparticle formulation has been clinically validated for safety in human mRNA vaccines and intravenous administration^28, 46^, considerable debate persists regarding the general bio-safety and potential adverse effects of LNPs in *in vivo* applications^14^. Here, we evaluated the in vivo biodistribution of the (LNP-CpG)+gE subunit vaccine candidate in mice and assessed its safety profile via whole-blood analysis in post-immunization rhesus macaques. In the murine fluorescence imaging study (Fig.S1), intense fluorescent signals for both the LNP vehicle and the gE antigen were preferentially enriched within intestinal tissue rather than the liver at 10 hours post-injection. This distinct trafficking profile suggests that the formulation may be preferentially drained to proximal lymphatic tissues, such as gut-associated lymphoid tissue (GALT), which is likely attributable to its optimal particle size. Concurrently, no distinct signal for the LNP-delivered CpG cargo was detected in the liver (Fig.S1B). We speculate that the absence of visible CpG fluorescence may be attributed to signal overlap, as short-wavelength red fluorophores such as Cy3 are readily absorbed by biological tissues, generating substantial autofluorescence background and exhibiting limited tissue penetration *in vivo* optical imaging. Nonetheless, this minimal hepatic accumulation circumscribes the risk of CpG- induced hepatotoxicity^47^. Following vaccine administration in rhesus macaques, no abnormal fluctuations were observed in body temperature or body weight (Fig.S3 a-c). Furthermore, comprehensive evaluations of CBC and serum biochemical profiles revealed that the fluctuations induced by the (LNP-CpG)+gE subunit vaccine were statistically indistinguishable from those elicited by the commercial Shingrix^TM^ (Fig.S2). Taken together, these multifaceted evaluations collectively demonstrate the outstanding biocompatibility and safety of the LNP-CpG platform for *in vivo* applications

In addition, LNP-CpG highlights its compelling viability as an alternative to the AS01B adjuvant in Shingrix^TM^. Developed by GSK, the AS01B and AS01E adjuvant systems are currently integrated into commercialized VZV and RSV subunit vaccines (Shingrix^TM^ and Arexvy^TM^, respectively). AS01B consists of DOPC, cholesterol, MPL (50 μg/dose), and QS21 (50 μg/dose), whereas AS01E contains the same active components at half the dosage. However, because QS21 must be harvested from the bark of South American Quillaja Saponaria trees, its supply is restricted by geographical and ecological limitations, ultimately driving up the production costs and market price of Shingrix^TM^. Conversely, LNP- CpG leverages components with highly mature manufacturing pipelines and fully synthetic, endlessly scalable CpG. Comparable to the performance of the AS01 platform, our LNP- CpG adjuvant demonstrated exceptional immunostimulatory capabilities for both VZV and RSV, whether formulated as single-antigen (Fig. 1, 2, 4) or combination vaccines (Fig. S4). Moreover, given that the therapeutic efficacy of the (LNP-CpG)+gE vaccine is significantly superior to that of Shingrix™ across multiple experimental dimensions, replacing GSK’s AS01 system with our LNP-CpG adjuvant holds substantial strategic potential.

A head-to-head comparison between the two HSV vaccine modalities revealed two distinctly contrasting outcomes (Fig. 5). The unadjuvanted LA-HSV vaccine, despite eliciting lower antibody titers and cellular immune responses, exhibited exceptional protective efficacy in the challenge model, both in preventing weight loss and in reducing viral loads. Consequently, while supplementing LA-HSV with LNP-CpG further boosted humoral and cellular immunity and lowered post-challenge viral titers, the additional gains were modest (Fig. 5h). This outcome is presumably attributable to the inherent immunological strengths of live-attenuated vaccines, their capacity to mirror natural infection and deliver continuous antigenic exposure. Conversely, the Alum+i-HSV performed poorly regarding both immune responses and protective efficacy, thereby rendering the advantage of LNP-CpG supplementation far more pronounced. Compared with the Alum+i-HSV cohort, the (LNP-CpG)+i-HSV formulation augmented immune response levels by several hundred-fold and achieved complete protection against post-challenge mortality and weight loss. In conclusion, although the LNP-CpG adjuvant consistently displays an outstanding ability to elevate immune responses across diverse vaccine platforms, the necessity of its implementation warrants multi-factorial evaluation.

There are several limitations in our study. Firstly, because an empty LNPs (eLNP), which consist of permanent cationic lipids, have been reported to be ineffective as adjuvants for subunit vaccines, they were excluded from the initial experimental design^9^. Therefore, the adjuvant potential of the eLNP formulation developed in this study, with ionizable lipids only, remains unassessed. Secondly, given literature reports that animal sex may influence immune responses^48, 49^, we stratified the animals by sex in the experiments for Figure S1 and Figure 2 to evaluate vaccine performance across sexes. However, no significant differences were observed between sexes in our studies, likely due to the limited sample size. Thirdly, in our current intracellular co-localization assays for LNP-CpG, the evaluation within plasmacytoid dendritic cells (pDCs) was overlooked. Given that pDCs are the primary immune cells expressing TLR9, MHC class II molecules and co-stimulatory molecules, have the potential for antigen presentation to CD4+ T cells^44, 50,51^, but with high difficulty in isolating from animals.

## Conclusion

In conclusion, this study demonstrates that packaging CpG ODN within lipid nanoparticles substantially amplifies its immunostimulatory potency. Notably, LNP-CpG exhibits versatile adjuvant activity that bridges distinct pathogens—including VZV, RSV, and HSV—and spans diverse vaccine platforms, encompassing subunit, inactivated, and attenuated formulations. Our findings highlight that LNP-CpG not only augments vaccine-induced immune responses but also imparts robust protective immunity against lethal viral challenges. Beyond these immediate applications, the LNP-CpG platform promises to unlock new paradigms for developing cross-protective, universal vaccines and next-generation adjuvant delivery technology.

## Methods

### 1. Materials and Reagents

Antigens (gE, Pre-F) were expressed in CHO system and purified by AtaGenix Laboratories Co., Ltd. (Wuhan, PR China). The CpG oligonucleotides (briefly called CpG, TGACTGTGAACGTTCGAGATGA) and Cy3-labeled CpG, possessing a complete phosphorothioate (PS) backbone, were synthesized by Sangon Biotech (Shanghai, China). Aluminium hydroxide gel adjuvant (10 mg/mL) was manufactured by Groda Denmark (Frederikssund, DK-3600, Denmark). Two vaccine candidates used in this study, live attenuated HSV-1 M4 (LA-HSV), and β-propiolactone inactivated HSV-1 ZW6 (GenBank: KX424525.1) (i-HSV), were prepared according to previously described protocol^52^, and kindly provided by Youchun Wang’s lab (IMB,CAMS. Kunming, China).

For cell experiment, all cell culture dishes/plates and centrifuge tubes were obtained from NEST Biotechnology Co., Ltd. (Wuxi, China). Cell culture medium and supplementary were purchased from VivaCell Biosciences (Shanghai, China). Cytochalasin D (CD, #B6645) and epoxomicin (EM, #A2606) were purchased from APExBIO Technology (Houston, TX, USA), bafilomycin A1 (BA1, #S1413) and concanamycin B (CB, #15502) was supplied by Selleckchem (Houston, TX, USA) and Cayman Chemical (Ann Arbor, MI, USA), respectively.

Pairs antibodies for sandwich ELISA and ELISpot in mouse study, including IL-2 (clone JES6-1A12 and JES6-5H4) and IFN-γ (clone AN-18 and XMG1.2), are purchased from Invitrogen (Carlsbad, CA, USA). ELISpot plates were supplied by Merck Millipore Ltd. Ireland. Tullagreen). For rhesus macaques’ study, commercial ELISpot kits from MabTech (Nacka Strand, Sweden) were used.

Fluorochrome-conjugated antibodies and related reagents utilized for flow cytometry were purchased from BioLegend (San Diego, CA, USA). Antibodies deployed in mouse experiments comprised PerCP/Cyanine5.5 anti-mouse CD4, FITC anti-mouse CD8a, APC anti-mouse IL-2, and PE anti-mouse IFN-γ. Meanwhile, the rhesus macaque panel consisted of anti-human CD4 (PerCP/Cyanine5.5), CD8 (FITC), IL-2 (APC), IFN-γ (PE), and TNF-α (Brilliant Violet 421™).

### 2. Animals and Ethics statement

C57bL/6J (female, six-week), Balb/C (female, six-week), and ICR (male and female, six-week) mice were purchased from Sipeifu (Beijing, China. Laboratory animal production license No. SCXK (jing) 2024-0001). Cotton rats were supplied and kept by Yishang Biotech Co., Ltd (Shanghai, China. Laboratory animal production license No. SCXK (hu)2022- 0011). All mice were kept in IMB,CAMS before use (Laboratory animal use license No. SYXK (dian) K2022-0006). All animal experiments were reviewed and approved by the Institutional Animal Care and Use Committee (IACUC) of IMB,CAMS, with committee number of: DWSP202410020, DWSP202503025, DWSP202411001, IACUC- 2024-CR-317, and DWSP202506021.

### 3. Preparation and characterization of LNPs

LNPs were synthesized using a Tofflon microfluidic formulation system (LNP-060L) equipped with its dedicated microfluidic chips (Nano Cartridge, Shanghai Tofflon Medical Packaging Material Co. Ltd), total flow rates were set at 12 mL/min. The ratio of liquid phase and lipid phase is 3:1. For liquid phase, CpG (120 μg/mL) alone, or with gE (60 μg/mL) mixed, were prepared in citrate buffer (pH 4.0). For lipid phase, (2-(2- hydroxyethoxy)ethyl)azanediyl)bis(hexane-6,1-diyl) bis(2-hexyldecanoate) (DHA-1), 1,2- distearoyl-sn-glycero-3-phosphocholine (DSPC), cholesterol, and 1,2-dimyristoyl-rac-glycero-3-methoxypolyethylene glycol-2000 (DMG-PEG2000) were, respectively, dissolved in ethanol at a molar ratio 40of 50:10:38.5:1.5, the total molecular mass is 12 mM. Immediately following nanoparticle synthesis, LNPs were dialyzed against an excess volume of Tris-Hcl (20 mM, pH 7.4) using Millipore Amicon Ultra-15 centrifugal filter units (100 kDa MWCO), and store at -20°C until use.

For analysis of particle size and polydispersity index (PDI), LNP samples were diluted 1:100 (v/v) in Tris-Hcl, and measured by Nano ZS Zetasizer (Malvern Instruments Corp., Malvern, UK). Each sample were repeated three times. A portion of the LNP sample was lysed by adding an equal volume of 10% Triton X-100, followed by the determination of the encapsulation efficiency of the cargo. CpG encapsulation efficiency of the LNPs was assessed using the RiboGreen™ ssDNA quantification assay kit; gE encapsulation efficiency was assessed using the BCA Protein Assay Kit, following the manufacturer’s instructions.

### 4. Cryogenic transmission electron microscopy (Cryo-TEM)

The TEM structural integrity of LNPs was assessed. By using a cryogenic sample preparation equipment (Vitrobot Mark IV, Thermo Fisher, USA), LNPs were loaded to grids (C-flat, R1.2/1.3 Au, 300 mesh, ZhongJingKeYi, China), which has been pre-hydrophilicity treated, and deposited for 45 s, followed by blotting with filter paper for 3 s. Imaging of the grids occurred using a CetaD/Falcon4 TEM (Thermo Fisher, USA) in brightfield mode at 200 kV.

### 5. Immunization and challenge

The immunization protocol for the animals is outlined in the corresponding schematic diagram (Fig.1d, 2a, 3a, 3e, S4a, 5a), following a two-dose schedule administered on days 0 and 28.

For the C57BL/6J mouse study, animals were randomly assigned to eight parallel groups (*n* = 6): (1) the gE-alone group, receiving 5 μg/dose of gE antigen; (2) the gE+CpG group, receiving 5 μg/dose of gE adjuvanted with free CpG (10 μg/dose); four distinct LNP formulation groups, comprising (3) LNP-encapsulated gE (7.5 μg/dose; LNP-gE), (4) LNP- gE supplemented with free CpG (10 μg/dose; (LNP-gE)+CpG), (5) LNP-encapsulated CpG (15 μg/dose) mixed with free gE (5 μg/dose; (LNP-CpG)+gE), and (6) co-encapsulated LNP containing both CpG (15 μg/dose) and gE (7.5 μg/dose); (7) the Shingrix^TM^ 1/10 group, where mice were injected with one-tenth of a human dose (1/10 hd) of the commercial vaccine; and (8) the control group, which received PBS injections as a blank control . At 14 days post-second immunization (on day 42), whole blood was collected from the mice to obtain immune sera, and the spleens were harvested for subsequent cellular analysis assays.

For the rhesus macaque study, animals were allocated into four parallel groups (*n* = 6), comprising a blank control group receiving PBS and three distinct vaccine groups: a 10-fold murine dose of (LNP-CpG)+gE, a hd of Ganwei^TM^, and a hd of Shingrix^TM^. Following both the prime and boost immunizations, the body weight and rectal temperature of the animals were closely monitored within the first 24 hours. Whole blood was collected to harvest immune sera at multiple pre-specified time points, including pre-immunization (day 0), 7 days post-prime (day 7), pre-boost (day 28), and 14 days post-boost (day 42). Concurrently, anti-coagulated blood samples were gathered on day 0, 28, and 42 to evaluate clinical chemistry, coagulation profiles, and hematological parameters. Finally, on day 42, peripheral blood mononuclear cells (PBMCs) were isolated from the whole blood for subsequent cellular analysis assays.

In the RSV-related studies, both Balb/C mice (*n* = 6) and cotton rats (*n* = 5) were utilized. For the murine study, the immunization regimen consisted of 2 μg/dose of Pre-F protein as the antigen, adjuvanted with either LNP-CpG (containing 14.36 μg/dose of CpG) or Alum (50 μg/dose). At 21 days post-second immunization (on day 49), immune sera were collected for antibody profiling. Subsequently, mice in the model control and immunized cohorts were intranasally challenged with RSV A2 (9.7 × 10⁵ PFU in 100 μL). At 5 days post-infection (5 dpi), lung tissues were harvested to quantify the viral load. In the cotton rat study, 5 μg/dose of Pre-F protein was deployed as the antigen to evaluate the Pre-F alone and the (LNP-CpG)+Pre-F formulations. Formalin-inactivated RSV (FI-RSV) vaccine was supplied by Yishang Biotech Co., Ltd (Shanghai, China. At 14 days post-second immunization (on day 42), immune sera were gathered from the cotton rats for antibody titration. Following this, each animal in the model and vaccine groups was intranasally inoculated with 1×10^5.5^ PFU of RSV A2. At 5 dpi, both lung and nasal turbinate tissues were harvested for further determination.

In the RSV+ VZV combined subunit vaccines study, Balb/C mice (*n* = 10) were immunized with Pre-F (5 μg/dose), gE (5 μg/dose), or both. Vaccines were adjuvanted with either LNP-CpG or Alum, as described above. The challenge viral dose and collected schedule were also same as above.

In the HSV-related studies, female Balb/C mice (3–4 weeks old, *n* = 21) were immunized intramuscularly twice at 4-week intervals (50 µL per injection). Two weeks after secondary immunization, partial mice were sacrificed for splenocytes assay (*n* = 6), remaining mice (*n* = 15) were challenged with wild-type HSV-2 (HG52) at a dose of 1.2 × 10^5^ PFU. Five days before challenge, mice were intramuscularly injected with medroxyprogesterone acetate (3 mg per mouse) to modulate their physiological cycle and increase their susceptibility to HSV-2. Vaginal swabs were collected at 0, 6, 8, 14 dpi, body weight, clinical symptoms and survival rates were monitored daily. At 7 dpi, 5 of 15 mice were sacrificed, vaginal tissues, spinals, and DRGs of mice were separated for viral loading detection.

### 6. Next-generation sequencing (NGS) analysis

At 24 hours post-prime immunization, whole blood was collected from the rhesus macaques for total RNA extraction, which was subsequently subjected to transcriptomic sequencing analysis on an Agilent platform (Agilent Technologies). The RNA was labeled with a Fluorescent Linear Amplification Kit according to manufacturer’s instructions. The quantity and labeling efficiency were verified before the samples were hybridized to whole-genome 8 × 60 k expression arrays, which were scanned at 5 μm using an Agilent scanner. Image analysis was performed with Feature Extraction software (version 11.5.1.1, Agilent Technologies) to generate raw microarray data.

### 7. Safety evaluation

For systemic safety evaluation, peripheral whole blood of rhesus macaques was collected on days 0, 14, and 28 post-immunization and divided into three aliquots: one was allowed to clot and centrifuged to obtain serum; one was collected into a heparin-anticoagulated tube as whole blood; and the third was collected into a heparin-anticoagulated tube, thoroughly mixed, and centrifuged to obtain plasma. Serum (400 μL) was analyzed for biochemical parameters (TBIL, ALT, AST, ALP, CREA, BUN, TC, and LDH) using an automatic veterinary chemistry analyzer (BS-200, Mindray, Shenzhen, China) with matching reagents (Mindray Bio-Medical Electronics Co., Ltd., Shenzhen, China). Heparin-anticoagulated whole blood (200 μL) was used to measure hematological parameters using an automated 5-part differential veterinary hematology analyzer (XT- 2000iV, Sysmex Corporation, Kobe, Japan). Plasma (400 μL) was used to evaluate coagulation parameters on an automated blood coagulation analyzer (CS-2400, Sysmex Corporation, Kobe, Japan) with matching reagents (Sysmex Jinan Medical Electronics Co., Ltd., Jinan, China). All assays were performed following the respective manufacturers’ instructions.

### 8. Antibody titers detection by indirect ELISA

The levels of antigen-specific antibodies in serum samples collected from immunized animals were determined by indirect ELISA. Briefly, antigens were dissolved in PBS and precoated in 96-well microplates, the final concentration of antigens (gE, Pre-F) are 2 μg/mL. After incubation overnight at 4 ℃, the plates were washed with PBST (0.05% (v/v) polysorbate 20 in PBS) 3 times. the plates were blocked with 5% (w/v) skim milk at 37 ◦C for 1 h, and serial diluted sera were added for another 1h. Goat anti-mouse HRP conjugated IgG (1:10,000), IgG2a (1:2000), and IgG1 (1:500) (Bio-Rad, Hercules, CA, USA) was used as the detection antibody. After addition of the mixed substrate 3,3,5,5- tetramethylbenzidine (TMB, Solarbio, Beijing, China) for 5 min, 1M sulfuric acid was added to stop the reaction. The absorbance at 450 nm was determined with a spectrophotometer (BioTek Instruments, Inc., Winooski, VT, USA). Antibody titers were defined by the end- point dilutions with a cutoff signal intensity of 2.1×OD450 of PBS control. Titers that lower than cutoff were set at 200 for calculations.

### 9. Live-virus neutralization assays

Detection of RSV-specific nAbs is essential for evaluating the immunogenicity of candidate RSV vaccines. In this study, an RSV cytopathic effect (CPE) neutralization assay was established using HEp-2 cells infected with either the RSV A2 or RSV B18537 strain as previously reported^53^. Briefly, RSV was co-incubated with serially diluted heat-inactivated test sera (starting at a 50-fold initial dilution, followed by 4-fold serial dilutions) in 96-well plates at 37°C for 1 h. Subsequently, 10,000 HEp-2 cells were seeded into each well. Following incubation at 37°C in a 5% CO_2_ incubator for 4–7 days, the cells were stained with a crystal violet solution for 30 min, after which viral CPE was observed and recorded. The numbers of CPE-positive and CPE-negative wells were accumulated, and the 50% inhibitory dilution (ID50) was determined as the serum dilution yielding a 50% reduction in CPE relative to the virus-only control wells. Finally, the neutralizing antibody titers (ID50) were calculated using the Reed–Muench method.

### 10. Cell isolation from animals and cell culture

Mouse spleens were dispersed with a 40 µm cell strainer, and single-cell splenocytes suspensions were harvested via centrifugation at 800× g for 5 min. Then, red blood cell lysis buffer (Serverbio, Wuhan, China) was added for another 5 min at room temperature. Then, cells were re-harvested via centrifugation at 800× g for 5 min. The splenocytes were counted and resuspended in Roswell Park Memorial Institute (RPMI) 1640 medium supplemented with 10% v/v fetal bovine serum (FBS) and 1% penicillin–streptomycin (complete RPMI 1640 medium, cRPMI) at a final concentration of 1×10^7^ cells/mL.

PBMCs were isolated from the whole blood of rhesus macaques using a commercial isolation kit (Solarbio, #P6720) according to the manufacturer’s instructions. Briefly, fresh anticoagulated whole blood was diluted with PBS and carefully layered over 3 mL of lymphocyte separation medium. After centrifugation at 500 × g for 20 min at room temperature, the lymphocyte layer was collected from the interface. The cells were washed twice with PBS and finally resuspended in cRPMI for subsequent experiments.

To generate bone-marrow-derived DCs (BMDCs), the femurs and the tibiae of C57BL/6J mice were removed, bone marrow cells were isolated and cultured the cells at 37 °C for 7 days with 20 ng/mL murine GM-CSF (PeproTech, Rocky Hill, NJ, USA). Cells were seeded and cultured in cRPMI with a density of 1 × 10^6^ cells/dish in petri dishes (Nunc, Roskilde, Denmark). These cells were stimulated with LNP-CpG (13.3 μg lipid with 0.5 μg CpG ODN/mL) at 37 °C for 24 h.

### 11. Cytokine detection by Sandwich ELISA

For the ELISA, each well of a 96-well plate was seeded with 100 µL of splenocytes at a final concentration of 1 × 10^6^ cells/well. Similarly, immunostimulants (gE, Pre-F, or inactivated HSV-1 antigen, 10 µg/mL) were then added to each well, along with an equivalent volume of PMA+ ionomycin (Dakewe Bioengineering Co., Ltd., Shenzhen, China) as a positive control. After incubation for 24 h at 37 ◦C, the supernatants were collected for cytokine level determination. Briefly, unconjugated anti-IL-2 (3 µg/mL) and anti-IFN-γ (4 µg/mL) antibodies, dissolved in PBS, were coated onto 96-well plates at 4 ◦C overnight. Subsequently, the plates were blocked with 1% (w/v) BSA in PBS at 37 ◦C for 1 h. Samples were added to each well and incubated for 3 h at room temperature. Biotin-conjugated antibodies against IL-2 and IFN-γ (2 µg/mL), dissolved in 1% BSA, were then added and incubated for another 1h, followed by the addition of HRP-conjugated streptavidin (1 µg/mL, Biolegend, Hercules, CA, USA), which was incubated for 30 min. TMB substrate and 2 mol/L sulfuric acid were sequentially added, and the absorbance at 450 nm was measured using a SYNERGY 4 microplate reader (BioTek Instruments, Inc., Winooski, VT, USA).

### 12. Cytokine spot detection by ELISpot

ELISpot plates were pre-coated with purified IL-2 and IFN-γ antibodies (2 µg/mL), and blocked with cRPMI before use. Splenocytes from mouse or PBMC from monkey were resuspended to a density of 1×10⁷ cells/mL in ELISpot-specific serum-free medium (Dakewe Biotech Co., Ltd., Shenzhen, China). The cells were then seeded at a density of 3 × 10⁵ cells per well and stimulated with antigens (final concentration of 20 μg/mL) for 20 h at 37°C in a 5% CO₂ humidified incubator. Following stimulation, plates were washed five times with PBS and incubated with biotin-conjugated anti-IFN-γ or anti-IL-2 antibodies for 2 h at room temperature. Subsequently, HRP-conjugated goat anti-mouse IgG secondary antibody (1:1000 dilution) was added and incubated for 1 h at room temperature. After five additional washes, TMB ELISpot substrate solution was applied to develop spots. The reaction was stopped by rinsing with water, and spots were quantified using an ImmunoSpot® S5 UV analyzer (Cellular Technology Co., Ltd.)

### 13. T cell differention by FC

A total of 1 × 10^6^ splenocytes (mouse) or PBMC (monkey) were incubated with 10 μg/ mL antigens at 37°C with 5% CO_2_ for 2 h, and 5 μg/mL brefeldin A was then added. Then the mixture was incubated overnight under the same conditions to block cytokine release. After washing with staining buffer, 100 μL of ZombieNIR™ was added to each vial, and the vials were incubated for 30 min. Then, 5 μg/mL anti-CD16/CD32 antibodies were added, and the cells were incubated at 4°C for 10 min to block nonspecific binding of Fc receptors. Thereafter, surface-staining antibodies (CD4, CD8) and intracellular-staining antibodies z were added subsequently (TNF-α, IFN-γ, IL-2), After staining for 30 min at 4°C, the cells were gated (forward and side scatter, FSC/SSC), and samples with more than 20,000 CD4^+^ T cells were analyzed with a CytoFLEX flow cytometer (Beckman, Indianapolis, IN, USA) and FlowJo V10 software (BD, Franklin Lakes, NJ, USA).

### 14. Tissue viral load detection by quantitative real-time PCR (qPCR)

qPCR analysis was performed using Premix Ex Taq (Probe qPCR, TaKaRa, Dalian, China) on a LightCycler 480 II platform (Roche, Indianapolis, IN, USA). To determine the RSV viral load, lung or nasal turbinate tissues were first ground, after which viral RNA was extracted using the Viral RNA Extraction Kit (QIAGEN, catalog no. 52906). After that, cDNA was synthesized with PrimeScript RT Master Mix (Takara, catalog no. RR036A). A standard curve was then plotted using the RSV N protein plasmid, and the viral load was calculated as the copies per milligram of lung tissue. qPCR targeting the RSV A2 strain N gene employed the following primers and probe: forward primer (5’- GGCAGTAGAGTTGAAGGGATTTC-3’), reverse primer (5’- TGCACACTAGCATGTCCTAAC-3’), and the TaqMan probe (5’-FAM- TATGAATGCCTATGGThsvGCAGGGCA-BHQ1-3′).

To determine the HSV viral load, DNA extraction from vaginal swabs and tissues was conducted using a MiniBEST Viral RNA/DNA Extraction Kit (TaKaRa, Dalian, China), according to the manufacturer’s protocol. The primers and TaqMan fluorescent probe targeting the HSV-2 gG gene were synthesized by Sangon Biotech (Shanghai, China) as follows: forward primer 5’-CGCTCTCGTAAATGCTTCCCT-3’, reverse primer 5’- TCTACCCACAACAGACCCACG-3’, and probe 5’-FAM- CGCGGAGACATTCGAGTACCAGATCG- BHQ1-3’.

### 15. H&E staining

Tissue samples were fixed in 10% formaldehyde for 24 h and dehydrated through a graded ethanol series, then embedded in paraffin wax, and sectioned at a thickness of 4 μm. Sections were mounted on microscopy slides (CITOGLAS, Jiangsu, China), stained with hematoxylin and eosin (H&E), and subsequently examined for histopathological changes using a Nikon eclipse TS2-FL optical microscope (Nikon, Tokyo, Japan).

### 16. Confocal imaging

Cy3 labeled CpG (^Cy3^CpG) were synthesized by Sangon, with Cy3 fluorescent group in the 3’- terminal of sequences. Protein gE (1 mg) were incubated with FITC (0.14 μM) in dark place for 60 min, then dialysis in sodium acetate solution for use. BMDCs were seeded into 8-well chamber slides in 500 μL cRPMI. Cells were incubated with 10 μg/mL ^FITC^gE- for 2 hours at 37℃. 0.5 μg/mL of LNP-^Cy3^CpG were added into wells. Meanwhile, 2 μg/mL of CD, 0.07 μg/mL of EM, 62.3 ng/mL of BA1 or 4.2 ng/mL of CB were added into corresponding wells. LNP-CpG and inhibitors were incubated with cells for 16 hours. Afterwards, cells were incubated with LysoTracker Deep Red (50 nM) for 1 hour and fixed with 4% formaldehyde. Images were obtained with Leica STELLARIS confocal microscopy.

### 17. *In vivo* image

For lipid labeled, 5 μM DiD perchlorate (dissolved in DMSO, 10 mg/mL) were added in the organic phase. gE (1 mg) were labeled with 800CW NHS ester as above described. ^DID^LNP-^Cy3^CpG were made follow the protocols described above, and 5 μg/dose ^800CW^gE were added. (^DID^LNP-^Cy3^CpG) +^800CW^gE (100 μL) were intramuscular injected into the right thigh of ICR mice. After injection, two female and two male mice were sacrificed for in vivo image in respective time point. Organs were dissected for observing the distribution of fluorescent signals within tissues.

### 18. Statistical analysis

Statistical analyses were performed using GraphPad Prism version 10.1.2 (GraphPad Software, La Jolla, CA, USA). Significant differences among groups were assessed by t’test, one-way analysis of variance (ANOVA), or two-way ANOVA. Significance was denoted as follows: *ns* for *p* > 0.05; * for *p* ≤ 0.05; ** for *p* ≤ 0.01; *** for *p* ≤ 0.001; and **** for *p* ≤ 0.0001. Illustration of animal models were formed by Biorender: https://app.biorender.com/profile/template/details/t-6a66c83d224364da058b70b2-lnp-cpg-deploy-the-self-adjuvant-role-of-mrna-vaccines

## Data availability

All data used during the study are available from the corresponding author upon request.

## Author contributions

Conceptualization, N.L. and C.L.; data curation, N.L.; formal analysis, N.L.; funding acquisition, N.L. and C.L.; investigation, N.L., H.C., X.Z., F.Y., C.L., Y.H., Y.B., and L.L.; methodology, N.L., H.C., X.Z., Z.H., S.F., and Q.L., project administration, C.L.; resources, Q.L. and S.W.; validation, C.L.; Y.H., Q.L., and S.F.; writing—original draft, N.L.; writing— review and editing, C.L. All authors have read and agreed to the published version of the manuscript.

## Acknowledgments

This research was supported by the National Key R&D Program of China (2022YFC2305700), Yunnan Fundamental Research Projects (202501AS070011 and 202601AT070221, 202401AT070148), the Special Biomedicine Projects of Yunnan Province, China (202602AS100008), Funds for the Training of High-Level Health Technical Personnel in Yunnan Province (D-2024054, H-2025005, and H-2025029), Funds for High-Level Scientific and Technological Talents Selection Special Project of Yunnan Province (202205AC160015), and The Scientific and Technological Innovation Fund of the Institute of Medical Biology, Chinese Academy of Medical Sciences (2026IMBCAMS002).

## Notes

### Competing Interest Statement

The authors have declared no competing interest.

