## Supplementary material for "LNP-CpG: deploy the self-adjuvant role of mRNA vaccines": Figure S

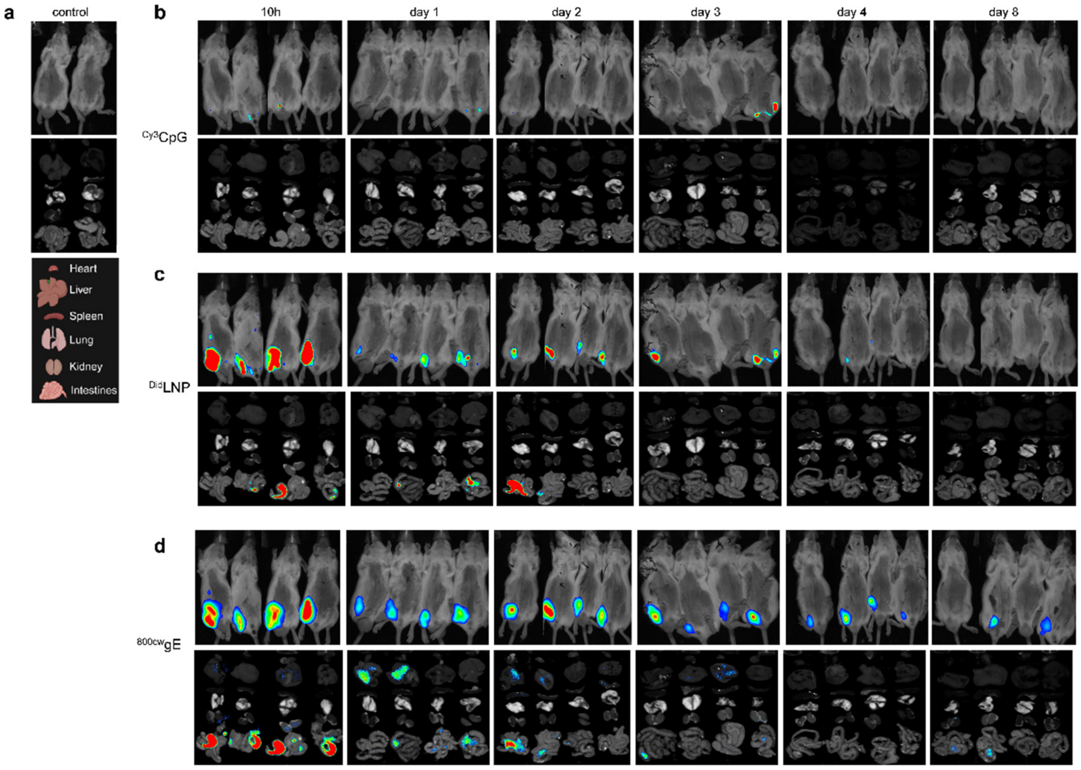


Figure S1. *In vivo* and *ex vivo* biodistribution kinetics of the (LNP-CpG)+gE vaccine candidate. Longitudinal *in vivo* fluorescence tracking and corresponding *ex vivo* organ imaging of mice following a single intramuscular injection of the formulated vaccine candidate over a series of predetermined time points (10 h, day 1, day 2, day 3, day 4, and day 8). Three distinct fluorescent channels were monitored simultaneously: (a) Imaging data of the control mice, harvested organs were arranged from top to bottom: heart, liver, spleen, lung, kidney, and intestines. (b) $^{Cy3}$CpG channel capturing the distribution and release kinetics of the nucleic acid cargo. (c) $^{Did}$LNP channel tracking the lipid nanoparticle vehicle. (d) $^{800CW}$gE channel demonstrating the localization of the external gE antigen. For each channel, the top row displays whole-body in vivo imaging, and the bottom row illustrates the *ex vivo* fluorescence profiles of harvested organs. Female and male mice are positioned on the left and right sides, respectively.

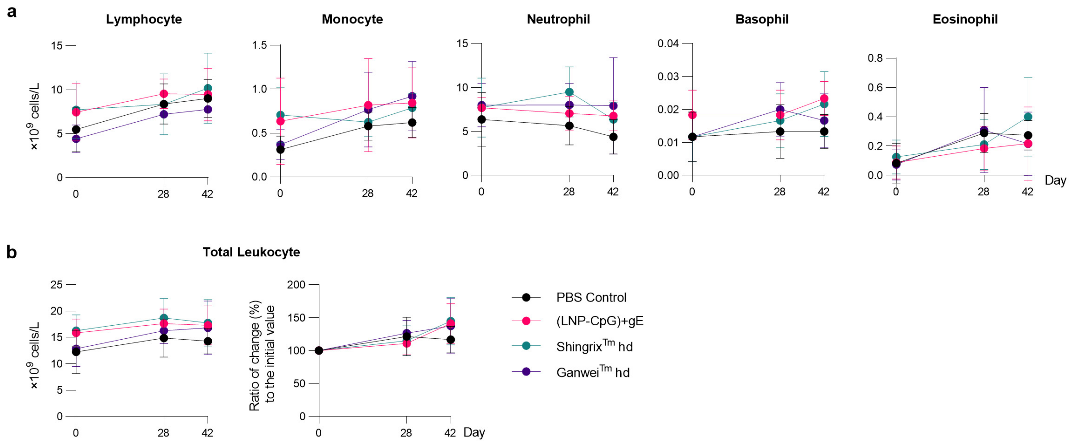


Figure S2. Measurement of complete blood counts in vaccinated rhesus macaques. Rhesus macaques were immunized i.m. with VZV vaccines, including (LNP-CpG)+gE, Ganwei$^{TM}$ hd, and Shingrix$^{TM}$ hd, with same volume PBS injected as control. (a) Complete blood counts, including lymphocyte, monocyte, neutrophil, basophil, and eosinophil, were measured from whole blood collected before vaccination and at day 0, 28 (before the second dose), and 42 (two weeks after second dose). (b) The total leukocyte count was calculated by summing all data points in (a). Additionally, the values on Day 0 were set to 100% to determine the percentage changes at subsequent time points after prime vaccination. hd: human dose. *n* = 6. Data are mean ± SD.

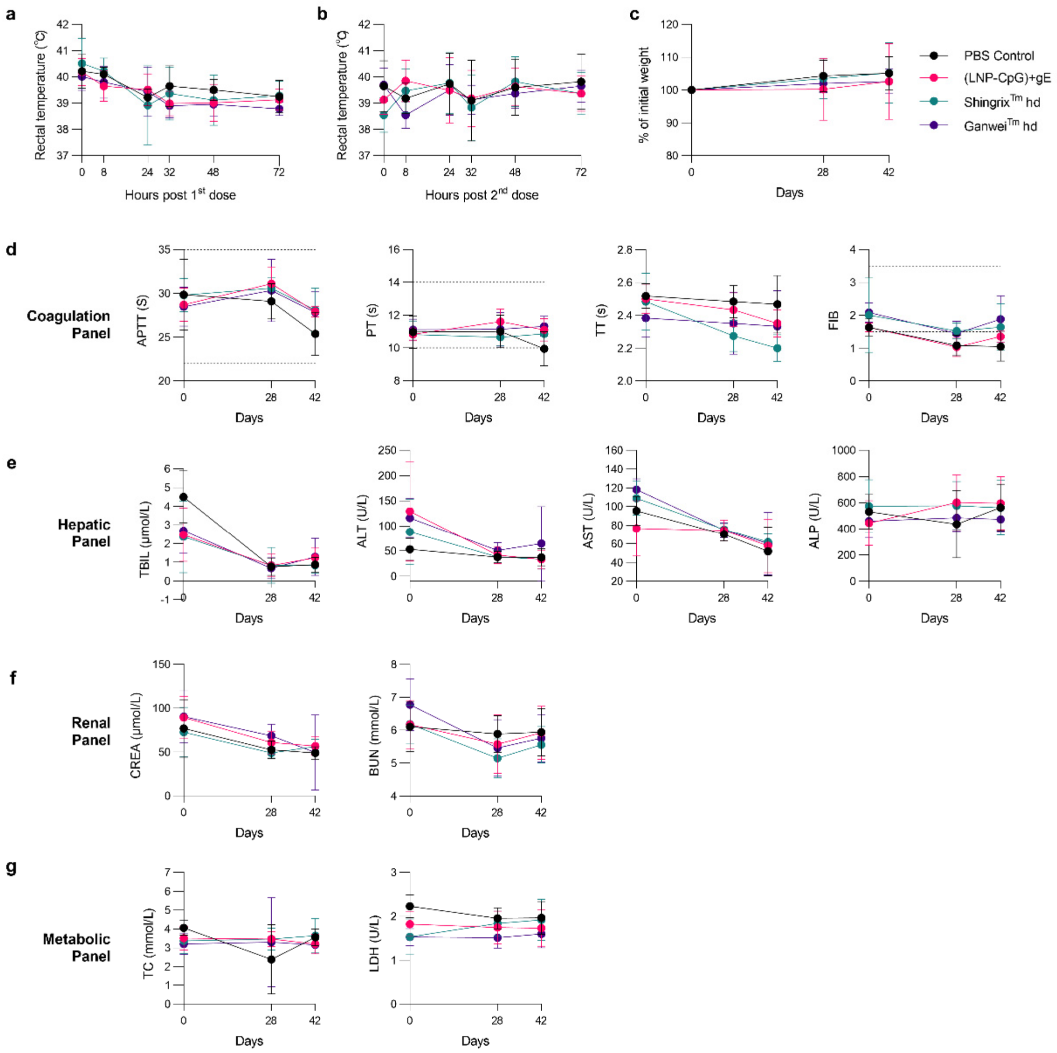


Figure S3. Safety profile after vaccines injected in rhesus macaques (*n* = 6). (a-b) Rectal temperature was monitored after first and second dose injected. c) % body weight change of initial. (d-g) Whole anticoagulated blood was collected on day 0, 28, and 42 for biochemical analyses. Indicator markers of coagulation panel, hepatic panel, renal panel, and metabolic panel were examined. Data are mean ± SD.

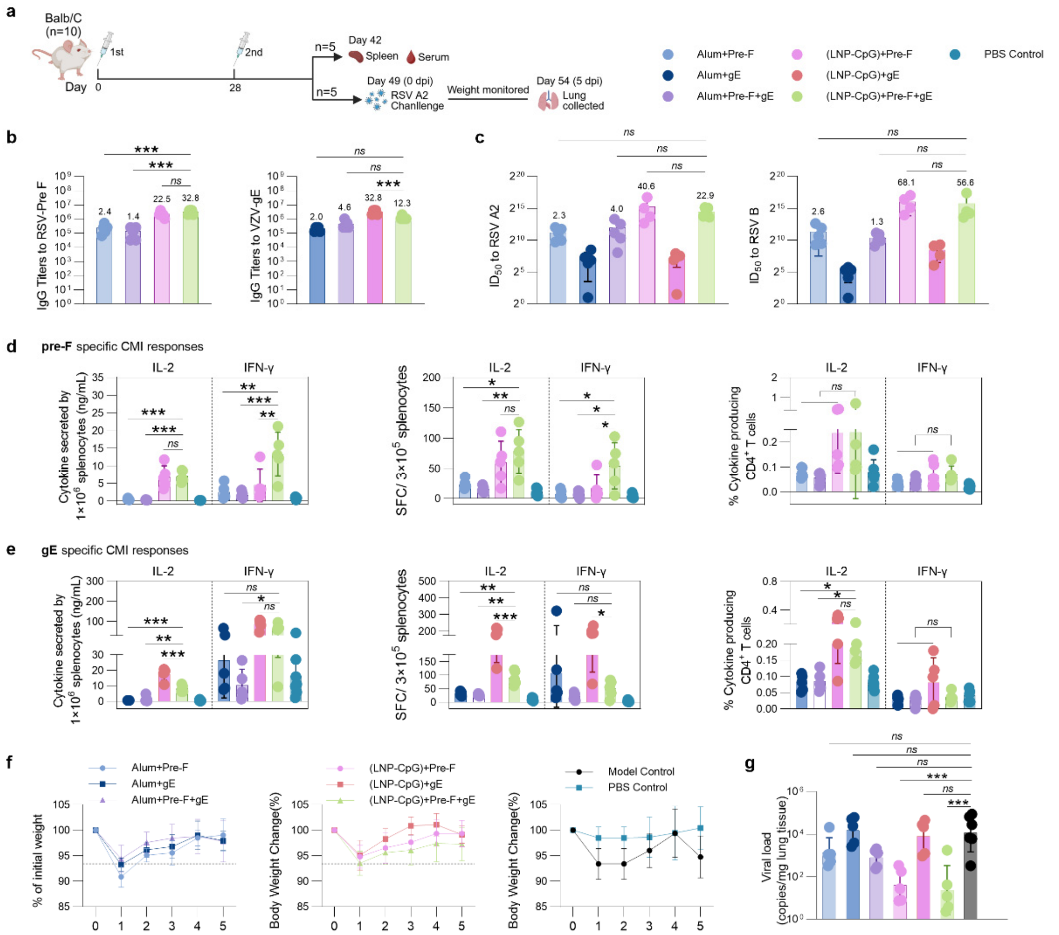


Figure S4. Immune profiling of Pre-F+gE combinate RSV+VZV subunit vaccines in mice. (a) Study design. Balb/C ($n$ = 10) were immunized 2 times at day 0 and 28. Splenocytes and immunized serum was collected at day 42 ($n$ = 5). Three weeks after second dose, mice were challenged by RSV A2. Body weight and survival rate were continue monitored for 5 days. Mice were sacrificed at 5 dpi; lung tissue were dissected for further test. (b) IgG titers specific to pre-F and gE. (c) nAbs titers specific to RSV A2 and RSV B strains were determined by an IC50 assay using HEp-2 cells. The recall responses of IL-2 and IFN-γ in splenocytes stimulated by pre-F (d) and gE (e) were evaluated using ELISA, ELISpot, and flow cytometry assays. (f) The proportional change in body weight compared with the 0 dpi baseline. Dashed lines represent the minimum value of the model control group for clarity. (g) Viral loads (copies/mg) of RSV A2 in lung tissues (5 dpi) were quantified by qPCR. In (b-e), data were analyzed by one-way ANOVA (using a Dunnett's multiple comparisons test with (LNP-CpG)+Pre-F+gE as a control column). In (g), data were analyzed by one-way ANOVA (using a Dunnett's multiple comparisons test with Model control as a control column). dpi: day post infection. Data were shown as mean ±SD. *, $p \leq 0.05$; **, $p \leq 0.01$; ***, $p \leq 0.001$; ****, $p \leq 0.0001$. Illustration in (a) created in BioRender.
